# The immunocytokine L19-TNF eradicates Sarcomas in combination with Chemotherapy agents or with immune check-point inhibitors

**DOI:** 10.1101/830406

**Authors:** Riccardo Corbellari, Lisa Nadal, Alessandra Villa, Dario Neri, Roberto De Luca

**Affiliations:** University of Trento, Italy, CiBIO (Department of Cellular, Computational and Integrative Biology), Via Sommarive 9, 38123 Povo (Trento); Philochem AG, Otelfingen, Switzerland; Department of Chemistry and Applied Biosciences, Swiss Federal Institute of Technology (ETH Zürich), Zurich, Switzerland

**Keywords:** Soft Tissue Sarcoma, Tumor targeting, Immunocytokine, TNF, Chemotherapy, Immune check-point inhibitors

## Abstract

Antibody-cytokine fusion proteins (also called “immunocytokines”) represent an emerging class of biopharmaceutical products, which are being considered for cancer immunotherapy. When used as single agents, pro-inflammatory immunocytokines are rarely capable of induce complete and durable cancer regression in mouse models and in patients. However, the combination treatment with conventional chemotherapy or with other immune-stimulatory agents typically increases the therapeutic efficacy of immunocytokines.

In this article, we describe combination treatments of a tumor-targeting antibody-cytokine fusion protein based on the L19 antibody (specific to a splice isoform of fibronectin) fused to murine tumor necrosis factor (TNF) with standard chemotherapy (dacarbazine, trabectedin or melphalan) or with an immune check-point inhibitor (anti-PD-1) in a BALB/c derived immunocompetent murine model of sarcoma (WEHI-164).

All combination treatments led to improved tumor remission compared to single agent treatments, suggesting that these combination partners may be suitable for further clinical development in sarcoma patients.

## Introduction

Sarcomas represent an heterogenous group of malignancies that can originate from soft tissue, cartilage, bone and other connective tissues. In particular, soft tissue sarcomas are tumors, which stem from non-epithelial extra skeletal tissue (e.g. muscle, fat, blood vessels and others). These cancer types are rare (< 1-2% of adult tumors) and they mainly affect young patients, however they represent 15% of all pediatric cancers [1,2].

For small and localized tumors (i.e. in the limbs) surgical resection (sometimes combined with radiation) represents a curative therapy, however postsurgical recurrence is observed in 50% of patients. This treatment option is often not applicable for patients with metastatic disease or with tumors with a different histology and in these cases, chemotherapy is administered as first-line treatment [3]. Doxorubicin represents the standard-of-care for sarcomas with a response rate of 12-30% [4]. Doxorubicin may be combined with dacarbazine or other agents, however these combination treatments typically do not increase the survival benefit compared to the monotherapy [5]. Moreover, they are usually associated with substantial toxicities.

Already in the late 19^th^ century, William B Coley observed a case of a sarcoma patient who experienced a spontaneous tumor regression as a consequence of an erysipelas infection. This finding was the basis for the development of the “Coley’s toxin”, a mixture of heat-inactivated bacteria, which was found to increase the overall survival of sarcoma patients, in spite of the development of high fever and chill [6,7]. It was later demonstrated that Coley’s toxin treatment induced a dramatic increase of TNF levels [8].

TNF is a homo-trimeric pro-inflammatory cytokine, mainly produced by macrophages, NK and T cells [9,10]. TNF is involved in inflammatory processes and as indicated by its name, it may display a potent anticancer activity against some malignancies, by promoting hemorrhagic necrosis and by inducing direct apoptosis of tumor cells [11–14]. Moreover, TNF may activate the immune system as a result of the increased inflammation [15–17].

The systemic administration of recombinant human TNF (Beromun^®^) in the clinic is limited by substantial side effects and the maximal tolerated dose (MTD) was found to be 300µg/m^2^ [18]. Beromun^®^ obtained marketing authorization only for locoregional treatments (i.e., for isolated limb perfusion procedure) in patients with soft-tissue sarcomas. The product is frequently administered in combination with melphalan [19].

The use of recombinant cytokines for cancer therapy may cause substantial toxicities already at low doses, therefore limiting their clinical use and preventing escalation to therapeutically active dose regimens [20–22]. To increase the therapeutic efficacy of cytokine payloads, the generation of fusion proteins with tumor-homing antibodies has been proposed. Antibody-cytokine fusion proteins (also termed “immunocytokines”) are capable of increasing the therapeutic index of the corresponding cytokine payload, as results of a selective localization at the site of disease [21–23].

We have previously described the immunocytokine L19-mTNF, consisting of the fully human antibody fragment L19 (specific to the alternatively spliced EDB domain of fibronectin, a marker of angiogenesis) [24,25] fused to murine TNF. The fusion protein was able to induce tumor regression in different murine cancer types either alone or in combination with chemotherapy or other immunocytokines [26–28].

The fully human L19-TNF (featuring human TNF) has been investigated in cancer patients (phase I and II completed) either as systemic administration, intralesional injection or in isolated limb perfusion settings [29–31]. Moreover, L19-TNF is currently being investigated in Phase III clinical trials for the treatment of Stage III B,C melanoma (EudraCT number: 2015-002549-72 and NCT03567889) in combination with L19-IL2 and of metastatic soft-tissue sarcoma (EudraCT number: 2016-003239-38 and NCT03420014) in combination with doxorubicin.

As previoulsy mentioned, melphalan, trabectedin and dacarbazine have been already considered for the treatment of sarcoma patients. For this reason, in this work we evaluated the combination of these chemotherapeutic agents with the immunocytokine L19-mTNF.

In the last decade, immune checkpoint inhibitors (e.g. anti CTLA-4, anti PD-1 and anti PD-L1) have shown encouraging clinical efficacy in the treatment of different malignancies [32–34]. However, not all cancer patients respond equally well to these drugs and combination strategies are currently being investigated in clinical trials aiming to improve therapeutic efficacy [35–37]. Moreover, the combination of immune checkpoint inhibitors with immunocytokines showed interesting results in mouse models of cancers [14,38]. Encouraged by these results, we now investigated the combination of an anti-PD-1 antibody with L19-mTNF for the treatment of sarcoma lesions.

L19-mTNF and anti PD-1 antibodies explicates their function in activating the immune system. For this reason, immunocompetent BALB/c mice were chosen as animal model. Mice models have become indispensable in pharmaceutical and biomedical research to verify the effectiveness of drugs and treatment regimens. These analysis may ensure safe and valuable data for a future clinical trial application [39].

### Aim of the work

The aim of this work was to investigate the combination of the immunocytokine L19-mTNF with chemotherapy agents (melphalan, trabectedin and dacarbazine) and with immune checkpoint inhibitors (anti PD-1) in a mouse model of sarcoma. The results of this work may provide a rationale and facilitate a future clinical investigation of these agents in sarcoma patients.

## Materials and Methods

### Cell lines and reagents

CHO cells and WEHI-164 were obtained from the ATCC. Cell lines were received between 2017 and 2019, expanded, and stored as cryopreserved aliquots in liquid nitrogen. Cells were grown according the supplier’s protocol and kept in culture for no longer than 14 passages. Authentication of the cell lines also including check of postfreeze viability, growth properties, and morphology, test for mycoplasma contamination, isoenzyme assay, and sterility test were performed by the cell bank before shipment.

The production and purification of L19-mTNF was performed as described before [26]. Commercially available dacarbazine (Dacin^®^, Lipomed AG), trabectedin (Yondelis^®^, PharmaMar AG), melphalan (Alkeran^®^, Aspen Pharma Schweiz GmbH) and the anti-PD-1 antibody (clone 29F.1A12, BioXCell, Catalog # BE0273) were purchased.

### Ethic statement

All animal experiments and maintenance were performed under a project license (license number 04/2018) granted by the Veterinäramt des Kantons Zürich, Switzerland, in compliance with the Swiss Animal Protection Act (TSchG) and the Swiss Animal Protection Ordinance (TSchV).

### Study design

Two different therapy experiments were performed with 24 and 30 BALB/c mice, respectively. One control group (mice injected only with saline) was used for each experiment, 5 groups were used as experimental. Mice were randomized into treatment groups according to their tumor volume (**Supplementary table 1 and 2**); measurements were taken by the same experimenter to minimize any subjective bias. Mice were divided in six groups of 4 and 5 animals (4 mice per group for the first reported therapy, 5 mice per group for the second therapy) per cage before the first drug injection. Blinded experiments were not performed.

### Experimental animals

A total of 54 female immunocompetent BALB/c mice, aged 8 weeks with an average weight of 20 g were used in this study. Mice were purchased from Janvier (Route du Genest, 53940 Le Genest-Saint-Isle, France).

Mice were raised in a pathogen-free environment with a relative humidity of 40-60% at a temperature between 18 and 26°C with daily cycles of 12 hours light/darkness according to guidelines (GV-SOLAS; FELASA). Mice were kept in an OHB Animal Facility in cages of maximum 5 mice, left for one-week acclimatization upon arrival and subsequently handled under sterile BL2 workbenches. Specialized personnel were responsible for their feeding; food and water were provided ad libitum. Mice were monitored daily (in the morning) in weight, tumor load, appearance (coat, posture, eyes and mouth moisture) and behavior (movements, attentiveness and social behavior).

Euthanasia criteria adopted were body weight loss > 15% and/or ulceration of the s.c. tumor and/or tumor diameter > 1500 mm and/or mice pain and discomfort. Mice were euthanized in CO_2_ chambers.

### Experimental procedure

Tumor cells were implanted subcutaneously in the flank of BALB/c mice using 5 × 10^6^ cells (WEHI-164) with 0.5 ml 29G insulin syringes (MicroFine™+, BD medical). Tumor volume was measured with a caliper and volume was calculated using the formula: Tumor Volume = (Length[mm]*Width [mm]*Width [mm])/2. When tumors reached a suitable volume (approx. 70-100 mm^3^), pharmacological treatment was performed.

L19-mTNF was dissolved in phosphate buffered saline (PBS), also used as negative control, sterile-filtered and administered at 0.1µg/g into the lateral tail vein three times every 48h. The intravenously route was chosen to achieve systemic administration of the bioactive payload. Dacarbazine (administered once intraperitoneally) and trabectedin (administered intravenously) were dissolved at 0.15 µg/g and injected four hours after the first L19-mTNF injection. Administration routes were chosen based on previously reported experiments [40,41].

The anti-PD-1 antibody was administered at 10 µg/g into the lateral tail vein three times every 48h alone or in combination with L19-mTNF. Melphalan was administered once intravenously into the lateral vein once at 4.5 µg/g. Also in this case the intravenously route was chosen to achieve systemic administration of the bioactive payload [28,38].

All injections were performed in the morning with 0.5 ml 30G insulin syringes (Omnican^®^ 50, Braun) in home cage for all mice groups. No anesthesia or analgesia were used for the injections.

### Experimental outcomes

The primary experimental outcome was the reduction in tumor volume of treated mice, indication of the effect of the administrated bioactive agents and drugs.

The survival end point was reached when the tumor diameter exceeded 1500 mm, body weight loss > 15% and/or ulceration of the s.c. tumor.

### Statistical method

Data were analyzed using Prism 7.0 (GraphPad Software, Inc.). Differences in tumor volume between therapeutic groups (until day 17, when n = 4 for the first therapy and until day 13, when n = 5 for the second therapy) were evaluated with the two-way ANOVA followed by Tukey post hoc test. P < 0.05 was considered statistically significant (*P < 0.05, **P < 0.01, ***P < 0.001, ****P < 0.0001).

## Results

In a first study, we compared the therapeutic activity of L19-mTNF, dacarbazine and trabectedin, either alone or in combination, in immunocompetent BALB/c mice bearing WEHI-164 lesions, a murine model of sarcoma, which is known to express EDB (the target antigen of the L19 antibody) [11,38] [**Figure 1**], according to the experimental scheme [**Figure 1A**].

**Figure 1.**
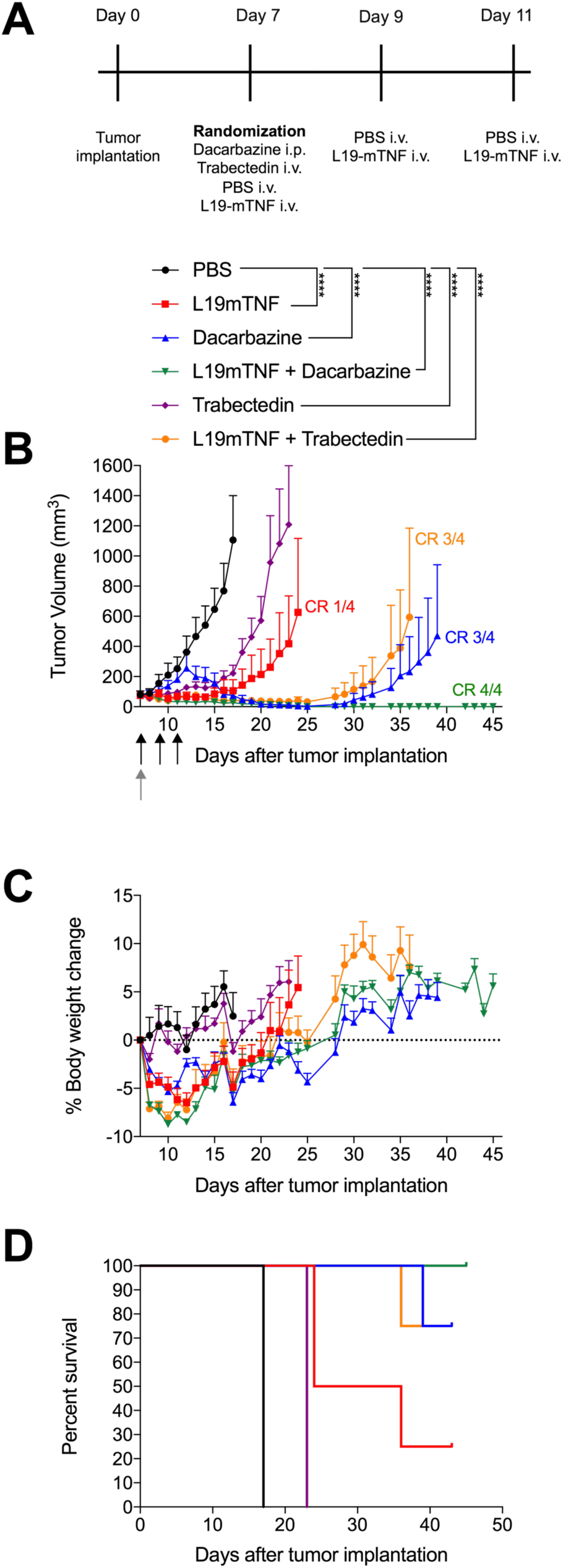
Therapeutic activity of L19-TNF in combination with chemotherapeutics (Dacarbazine and Trabectedin) in BALB/c mice bearing WEHI-164 tumors. (**A**) Experimental scheme. (**B**) Treatment started when tumors reached a volume between 70-100 mm^3^, mice were injected three times intravenously (black arrows) every 48 hours with either PBS or L19-TNF. Dacarbazine (injected intraperitoneally) and Trabectedin (injected intravenously) were administered once (gray arrow), four hours after the first L19-TNF injection (combination group). Data represent mean tumor volume ± SEM, n = 4 mice per group, CR = complete response. (**C**) Body weight monitoring. Weight change during the therapy of subcutaneous WEHI-164 tumors. Data represent mean % weight (± SEM), n = 4 mice per group. (**D**) Survival curves.

As expected, the treatment with saline was not able to induce cancer regression. When used as single agents, L19-mTNF and trabectedin exhibited a mild tumor growth retardation compared to the saline group. In addition, 1 mouse out of 4 treated with L19-mTNF showed a complete response. However, the combination therapy with these two agents led to an enhanced antitumor effect (3 out of 4 complete response). Dacarbazine administered as a single agent was capable of eradicate tumors in 3 out of 4 mice. The therapeutic efficacy was improved when administered in combination with L19-mTNF with (4 out of 4 complete response) [**Figure 1B, D**]. All treatments were well tolerated as evidenced by the comparison of body weight profiles. Only a transient toxicity was observed upon the first pharmacological treatment with the two combination treatments (L19-mTNF with either dacarbazine or trabectedin). However, all treated mice recovered the body weight loss in few days [**Figure 1C**].

In a second experiment, we compared the therapeutic activity of L19-mTNF in combination with melphalan or with an immune check-point inhibitor (anti-PD-1) in the same murine sarcoma model (WEHI-164) [**Figure 2**], according to the experimental scheme [**Figure 2A**]. The treatment with saline and melphalan alone was not able to induce cancer regression, by contrast, when used as single agent, L19mTNF was able to cure 1 mouse out of 5. Treatment with anti-PD1 alone induced cancer eradication in 2 out of 5 mice. The therapeutic activity of L19-mTNF was potentiated when used in combination with both melphalan (3 out of 5 complete response) or with an anti-PD1 antibody (4 out of 5 complete response) [**Figure 2B, D**]. All the treatments were well tolerated as evidenced by the comparison of body weight profiles [**Figure 2C**].

**Figure 2.**
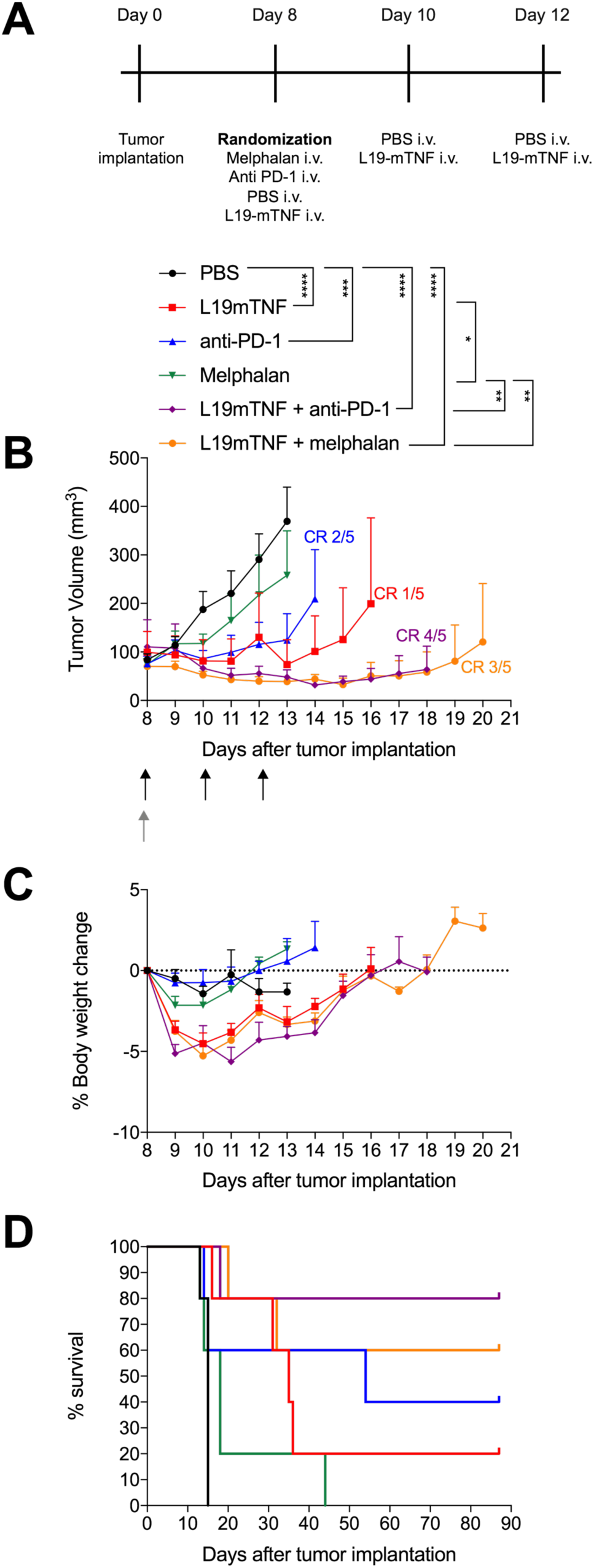
Therapeutic activity of L19-TNF in combination with immune-checkpoint inhibitor (anti-PD1) or chemotherapeutics (Melphalan) in BALB/c mice bearing WEHI-164 tumors. (**A**) Experimental scheme. (**B**) Treatment started when tumors reached a volume between 70-100 mm^3^, mice were injected three times intravenously (black arrows) every 48 hours with either PBS, L19-TNF, anti-PD-1 or the combination of L19-TNF plus anti-PD1. Melphalan were also administered intravenously once (gray arrow), both as a monotherapy and in combination with L19-TNF. Data represent mean tumor volume ± SEM, n = 5 mice per group, CR = complete response. (**C**) Body weight monitoring. Weight change during the therapy of subcutaneous WEHI-164 tumors. Data represent mean % weight (± SEM), n = 5 mice per group. (**D**) Survival curves.

## Discussion

We have now reported a pre-clinical evaluation of combination therapies with the fusion protein L19-mTNF. BALB/c mice bearing WEHI-164 lesions were used for this purpose. In this study, the 3Rs principle was constantly adopted checking mice health and comfort and minimizing pain or distress of the animals during the treatment administration.

During these studies, we observed a synergistic effect of L19-mTNF when used in combination with chemotherapeutic agents (trabectedin, dacarbazine and melphalan) as well as with an anti-PD1 antibody.

TNF is a proinflammatory cytokine able to activate the immune system and exert potent antitumor effects both in animal models and in patients [42]. Low-dose of TNF can indeed mediate the activation of CD8 T^+^ cells and their infiltration in the tumor microenvironment. TNF can also alter the vascular bed improving its permeation and causing endothelial activation as well as massive hemorrhagic necrosis enhancing antitumor effects [43].

In this study, the immunocytokine L19-mTNF, specific to EDB, increases the activity of lymphocytes into the tumor mass with no toxicity issue. Furthermore, previous studies revealed the ability of TNF to rapidly eradicate sarcoma tumor cells due to their sensibility towards TNF [11–13,44].

A possible explanation of the success of L19-mTNF when used both in combination and as single agent, may be amenable to the expression of immunodominant AH1 (SPSYVYHQF), a peptide derived from the gp70 envelope protein of murine leukemia virus. AH1 is endogenous in BALB/c genome and displayed on murine tumors (e.g. WEHI-164, C51 and CT26 tumor cells). SPSYVYHQF sequence is then recognized by CD8^+^ T cells previously expanded by the administration of L19-mTNF [12,44].

When used in combination, L19-mTNF potentiates the immunomodulatory effects of the chemotherapeutic agents.

Trabectedin is an immunomodulatory agent capable of inducing apoptosis of sarcoma cells through the inhibition of the production of IL-6 and CCL2 and the activation of TRAIL pathway, toxic for human monocytes [45].

It is reported that the Isolated Limb Perfusion with TNF-α in combination with melphalan for the treatment of sarcoma reached high response rate in multicenter trails. Melphalan showed a synergistic effect with TNF directing cytotoxic of tumor associated vasculature and tumor cells [46]. Results of melanoma patients treated with melphalan showed an expansion of CD8^+^ T cells, with enhanced levels of cytotoxic IFN-γ and granzyme B and expression of immune related markers such as MHC class I and Hsp70 [47].

Similarly, dacarbazine exerts antitumoral effect through the stimulation of the immune system. It acts inducing local activation of NK cells and CD8^+^ T cells. It is indeed already an established treatment for sarcoma patients and in this study, we confirmed its potency as single agent [48,49]. When dacarbazine was used in combination with L19-mTNF, 4/4 mice were cured thus confirming a potential synergistic effect between the two agents.

Immune checkpoint inhibitors are rapidly gaining acceptance as broad-spectrum agents for many cancer indications. The immunomodulatory effect of Nivolumab (anti-PD1 antibody) was showed by Choueiri’s work in 2016, where they demonstrated the reverse T-cell exhaustion in the tumor microenvironment at several doses. Nivolumab is able to increase both tumor-associated lymphocyte markers (CD3^+^ and CD8^+^) as the expression of hallmarks of Th1 inflammatory response (e.g. ICOS, IFN-γ, granzymes and perforin). It was also demonstrated a significant increase in the expansion of NK cells, suggesting that Nivolumab can enhance the T-cell-mediated immune activity [50].

We proved that L19-mTNF efficiently synergized with PD-1 blockade, thus providing evidence for a further clinical investigation in sarcoma patients.

## Conclusions

In this work, we presented a pre-clinical evaluation of combination modalities for the treatment of sarcoma lesions. The immunocytokine L19-mTNF was able to synergize with chemotherapeutic agents, which are already being considered for the treatment of sarcoma patients. Moreover, L19-mTNF was able to potentiate the anti-cancer activity of an anti-PD-1 antibody. The fully-human analog of L19-mTNF (L19-TNF), which is currently being investigated in phase I clinical trials in combination with doxorubicin, may also be considered in combination with other chemotherapeutic agents or with immune check-point inhibitors for the treatment of sarcoma patients. This study may also provide pre-clinical evidences for the upcoming phase I clinical trial of L19-TNF in pretreated Soft Tissue Sarcoma in combination with dacarbazine foreseen for 2020.

## Acknowledgments

We would like to thank Dr. Teresa Hemmerle for helpful discussions and we gratefully acknowledge Alessandro Pedrioli for his help with experimental procedures.

