## Supplementary Tables for "The immunocytokine L19-TNF eradicates Sarcomas in combination with Chemotherapy agents or with immune check-point inhibitors" (from another version of this preprint: version 2, 2019-12-11)

**Supplementary Information**

**Table 1**

| | PBS | L19-mTNF | dacarbazine | trabectedin | L19-mTNF + dacarbazine | L19-mTNF + trabectedin |
|---|---|---|---|---|---|---|
| Average ($mm^3$) | 80.6 | 75.9 | 75.5 | 72 | 73.2 | 75 |
| Standard Dev. ($mm^3$) | 46.8 | 41.1 | 37.8 | 29.8 | 33.8 | 24.6 |

**Table 1** Tumor volume values and Standard Deviation of mice at randomization day used in the therapy with L19-mTNF in combination with trabectedin and dacarbazine.

**Table 2**

| | PBS | L19-mTNF | a-PD1 | melphalan | L19-mTNF + a-PD1 | L19-mTNF + melphalan |
|---|---|---|---|---|---|---|
| Average ($mm^3$) | 84.1 | 99.4 | 75.8 | 73.5 | 110.7 | 70.4 |
| Standard Dev. ($mm^3$) | 27.1 | 96.2 | 35.7 | 27.8 | 124.8 | 29.7 |

**Table 2** Tumor volume values and Standard Deviation of mice at randomization day used in the therapy with L19-mTNF in combination with a-PD1 and melphalan.
